# Nonsynonymous Mutations in Intellectual Disability and Autism Spectrum Disorder Gene PTCHD1 Disrupt N-Glycosylation and Reduce Protein Stability

**DOI:** 10.1101/2022.09.23.509248

**Authors:** Connie T. Y. Xie, Stephen F. Pastore, John B. Vincent, Paul W. Frankland, Paul A. Hamel

## Abstract

*PTCHD1* has been implicated in Autism Spectrum Disorders (ASD) and/or intellectual disability, where copy number variant losses or loss-of-function coding mutations segregate with disease in an X-linked recessive fashion. Missense variants of *PTCHD1* have also been reported in patients. However, the significance of these mutations remains undetermined since the activities, subcellular localization and regulation of the PTCHD1 protein are currently unknown. This paucity of data concerning PTCHD1 prevents the effective evaluation of sequence variants identified during diagnostic screening. Here, we characterize PTCHD1 protein binding partners, extending previously reported interactions with postsynaptic scaffolding protein, SAP102. Six rare missense variants of PTCHD1 were also identified from patients with neurodevelopmental disorders. After modelling these variants on a hypothetical three-dimensional structure of PTCHD1, based on the solved structure of NPC1, PTCHD1 variants harboring these mutations were assessed for protein stability, post-translational processing and protein trafficking. We show here that wild-type PTCHD1 post-translational modification includes complex *N*-glycosylation and that specific mutant proteins disrupt normal *N*-link glycosylation processing. However, regardless of their processing, these mutants still localized to PSD95-containing dendritic processes and remained competent for complexing SAP102.

## Introduction

High throughput and increasingly precise genomic approaches have identified myriad genetic loci involved in Autism Spectrum Disorder (ASD) (1, 2). The biological pleiotropy of these defined loci, using cytogenetics, linkage analysis, whole-genome linkage or association as well as whole-genome or exome sequencing, underlines the complexity of ASD (3). The ASD-associated gene at Xp22.11, *PTCHD1*, was identified by several groups (4–8) using distinct approaches including one that indicated that *PTCHD1-related* mutations may occur in approximately 1% of individuals with ASD (9).

*PTCHD1* encodes an 888 amino acid protein that is structurally similar to the class of resistance-nodulation-cell division (RND) superfamily of transporters (for review see (10)) as well as two cholesterol transporters related to Niemann-Pick syndrome type C protein, NPC1 (11–13). While related to the receptors of the Hedgehog (Hh)-ligands, Ptch1 and Ptch2 (14–16), we and others have not yet been able to show that PTCHD1 plays a regulatory role in the Hh-pathway (17, 18) and there is a current lack of evidence showing that PTCHD1 directly binds to or facilitates cholesterol fluxes. Regardless, PTCHD1 encodes a protein predicted to harbor two “Ptch1-domains”. These modules, that are juxtaposed in the membrane, are defined by 5 transmembrane a-helices flanked by luminal and cytoplasmic regions. The luminal domains exhibit sequence similarities to the analogous regions in NPC1 (12, 19) and Ptch1 (20–22) that suggest they may have highly similar three-dimensional structures. However, like all other members of this class of transmembrane proteins, the cytoplasmic regions of PTCHD1 are unrelated to those in, for example, any of the Ptch-proteins or NPC1 (14). In the case of PTCHD1, the last four amino acids at its C-terminus encodes a unique motif predicted to bind PDZ-domain-containing factors (18). Using a yeast two-hybrid screen, this motif was used previously to isolate PSD95 and SAP102, proteins localized to dendritic spines in the post-synaptic region where a large number of factors involved in synaptic transmission are organized (18). Indeed, a GST-fusion protein encoding the C-terminus of PTCHD1 binds PSD95 although localization of PTCHD1 to dendritic spines did not appear to be dependent on the PDZ-binding region in its C-terminus, consistent with PTCHD1 transport to dendritic spines may being mediated by distinct mechanisms and regions of the protein.

We report here the identification and characterized of a series of point mutations in PTCHD1 derived from patients with ASD or other neurodevelopmental disorder. We show further that for a number of mutations in PTCHD1, these align with sequences crucial for the complexing with cholesterol in structurally-related protein, Ptch1. Despite these specific mutations altering the processing of the newly synthesized protein as well as the protein stability of PTCHD1, they do not produce defects on its ability localize to structures containing PSD95.

## Results

### Characterization of PTCHD1 mutants

We identified PTCHD1 missense variants from clinical studies where variants were identified in male individuals presenting with a neuro-developmental disorder and was not present in the control database (gnomAD:gnomad.broadinstitute.org; from >182,000 exomes + genomes sequenced). For these variants, we used two methods: **i)** the Condel missense prediction meta-algorithm, which combines predictions from five algorithms (23) and **ii)** the Combined Annotation-Dependent Depletion (CADD) (24) to predict whether the substitutions are likely to be deleterious.

Based on these analyses, we selected a subset of variants (**Figure 1A**) i) predicted by both methods to be deleterious, ii) spanning the PTCHD1 protein and iii) representing mutations in distinct structural regions. These regions included the first and second luminal domains, “Loop 1” and “Loop 2”, and the two transmembrane modules that produce the 3D structures resembling sterol sensing domains (“Ptch-domains”), SSD1 and SSD2 (**Figure 1B-D**). For comparison purposes we also included a nonsense mutation (ΔITTV) that disrupts the predicted C-terminal PDZ-binding motif, although this deletion does not correspond with a known clinically-reported mutation.

**Figure 1.**
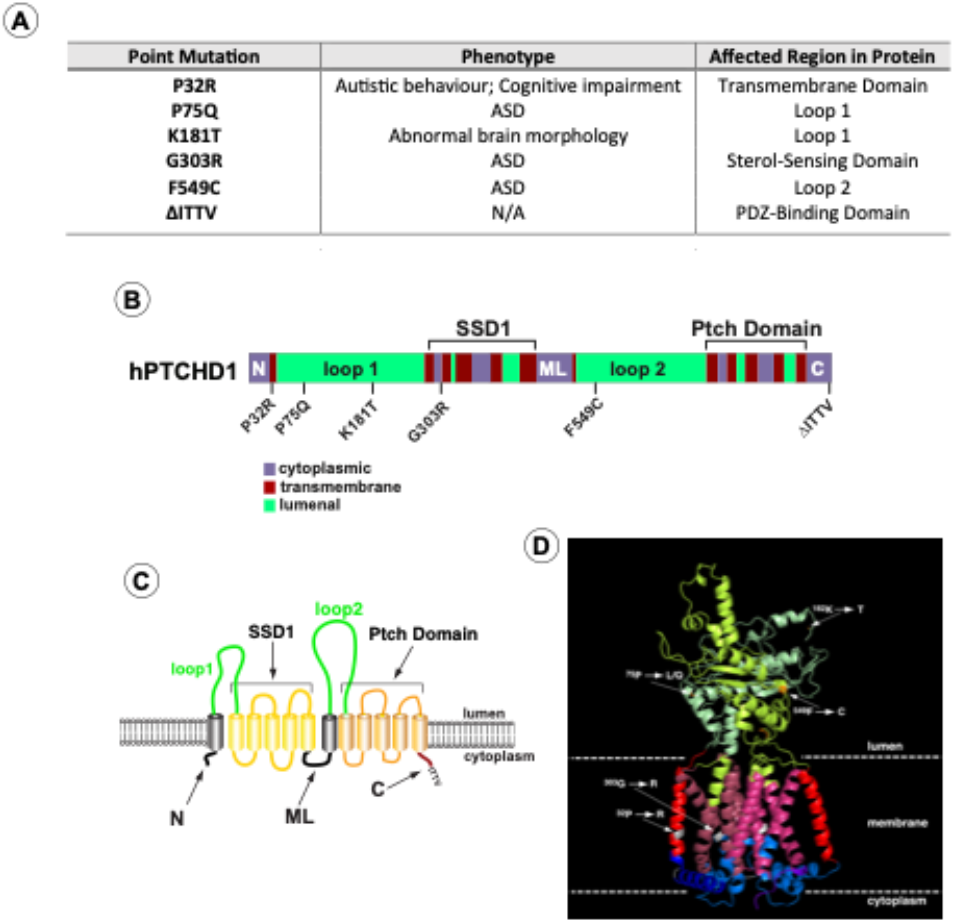
PTCHD1 graphics depicting point mutations and predicted structure. **A)** Summary of point mutants in PTCHD1 from clinical studies (see also Appendix 1) **B)** Linear schematic of PTCHD1 indicating structural domains and locations of point mutations. **C)** Cartoon of PTCHD1 illustrating the predicted topological orientation of specific regions in the membrane. **D)** A predicted 3D structure of PTCHD1 based on the resolved cryo-EM structures of NPC1 and Ptch1. Locations of the point mutations in the hypothetical structure are indicated. Note that the cytoplasmic domains cannot be resolved for this class of proteins and are, therefore, absent.

As the linear cartoon in **Figure 1B** illustrates, two of these mutations (P32R and G303R) produce amino acid changes in a-helical regions that are predicted to traverse the membrane. In contrast, P75Q, K181T and F549C alter amino acids in the luminal domains. Using Phyre^2^ to align the sequence for PTCHD1 with that of human NPC1 and Ptch1, whose 3D structures have been determined previously (11, 20–22, 25, 26), we used PyMol to generate a hypothetical three-dimensional structure for PTCHD1 (**Figure 1D**). For all regions except the N, ML and C cytoplasmic domains, this predicted model is nearly identical to a recent 3D model predicted using AlphaFold (27) (see Appendix 1). As illustrated on the hypothetical model in Fig. 1D, the P32R and G303R mutations are expected to disrupt helical structures in the first two highly conserved alpha-helices in the T-class of SSD-containing proteins (28). G303R specifically affects the first helix in the SSD1-like domain, being the integrity of this domain is essential for the activities of both NPC1 (29) and Ptch1 (30–33).

We first sought to determine differences in the relative levels of protein expression due to mutations on PTCHD1. GFP cassette was fused N-terminal end of PTCHD1 to create the ^GFP^PTCHD1 construct, which was then cloned in frame with the GFP cassette through the P2A site in the pULTRA lentiviral vector. This arrangement produces a single transcript encoding a GFP-P2A-^GFP^PTCHD1 fusion protein. The P2A site facilitates proteolytic cleavage between the GFP and ^GFP^PTCHD1 proteins allowing direct comparison of their expression. As **Figure 2A** demonstrates that in HEK293 cells, the level of transcripts encoding the GFP-P2A-^GFP^PTCHD1 fusion protein for all mutants of ^GFP^PTCHD1 was essentially identical, differing by less than 10%. In contrast to these levels of message, **Figure 2B** illustrates that the levels of protein for these mutants varied considerably. Quantification of the levels of ^GFP^PTCHD1 normalized to those of the GFP alone and β-tubulin, confirmed that several of the point mutations were expressed at considerably lower levels of protein. (**Figure 2C**). Thus, despite their essentially identical levels of mRNA expression, the resultant proteins show considerable heterogeneity in their level of protein expression.

**Figure 2.**
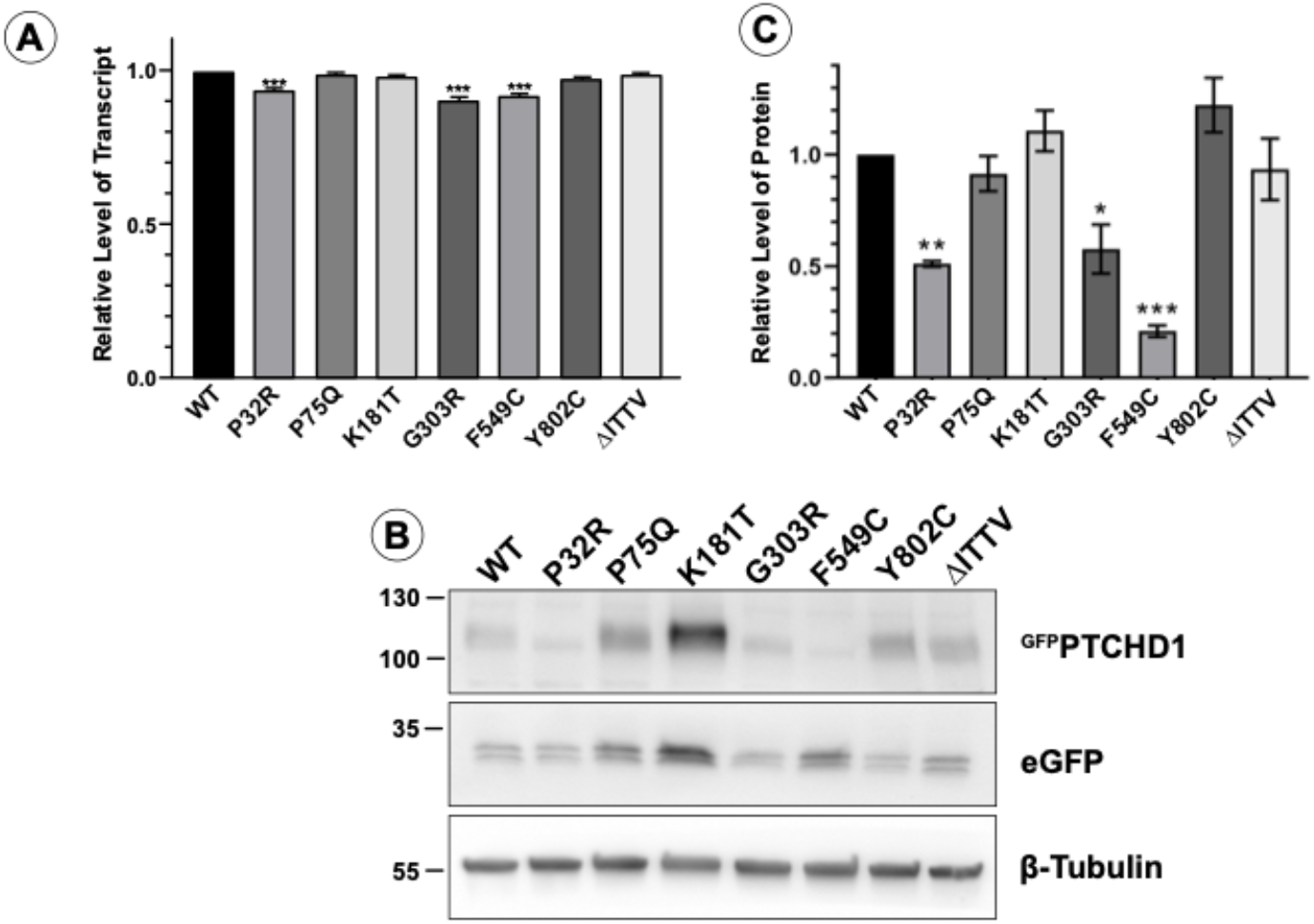
PTCHD1 mutants in stably-expressed cells vary in protein expression levels. **A)** Relative levels of stably-expressed ^GFP^PTCHD1 transcript in bulk cultures of HEK293 cells. **B)** Representative western blot ^GFP^PTCHD1, eGFP and β–Tubulin for stable lines expressing the PTCHD1 mutants **C)** Quantification of relative levels of expression of individual mutants stably-expressed in HEK293 cells. Data displayed as mean ± SEM, n=3 independent experiments. Data was analyzed using one-way ANOVA followed by Dunnett’s multiple comparisons test of means to the control (WT) using t-test. * p ≤ .05, ** p ≤ .01, *** p ≤ .001.

### Distinct processing of PTCHD1 mutants

Close inspection of the protein bands seen of ^GFP^PTCHD1 in **Figure 2B** shows that these mutant proteins have distinct patterns of migration under denaturing conditions in SDS-PAGE gels. Given the variation in expression patterns and levels of these mutant proteins, we characterized their post-translational processing, protein stability and subcellular localization. **Figure 3A** illustrates that wildtype ^GFP^PTCHD1 is processed to complex *N*-linked glycosylated forms. Here, the migration of the bulk of ^GFP^PTCHD1 is unaffected by treatment with EndoH, indicating processing of the *N*-linked glycosylated moieties to more mature, complex structures for the wild-type protein. **Figure 3B** illustrates mutants with contrasting patterns of processing. In the case of the mutant protein that deletes the PDZ-binding motif, ΔITTV, a pattern of slower migrating species similar to the WT ^GFP^PTCHD1 is evident, the slowest migrating (upper arrow) also resistant to the EndoH. In contrast, the F549C point mutant exhibits only two slower-migrating species, both of which are susceptible to EndoH, consistent with this protein not being processed through Golgi-dependent transport pathways that generate mature *N*-link glycosylated protein. Using the same analysis (**Figure 3C**), the point mutants P75Q and K181T both gave rise to slow migrating, EndoH-resistent bands similar to WT ^GFP^PTCHD1. In contrast, P32R and G303R did not exhibit mature glycosylated forms since all forms are susceptible to EndoH activity. Thus, these different point mutants distinctly alter the apparent post-translational processing of PTCHD1.

**Figure 3.**
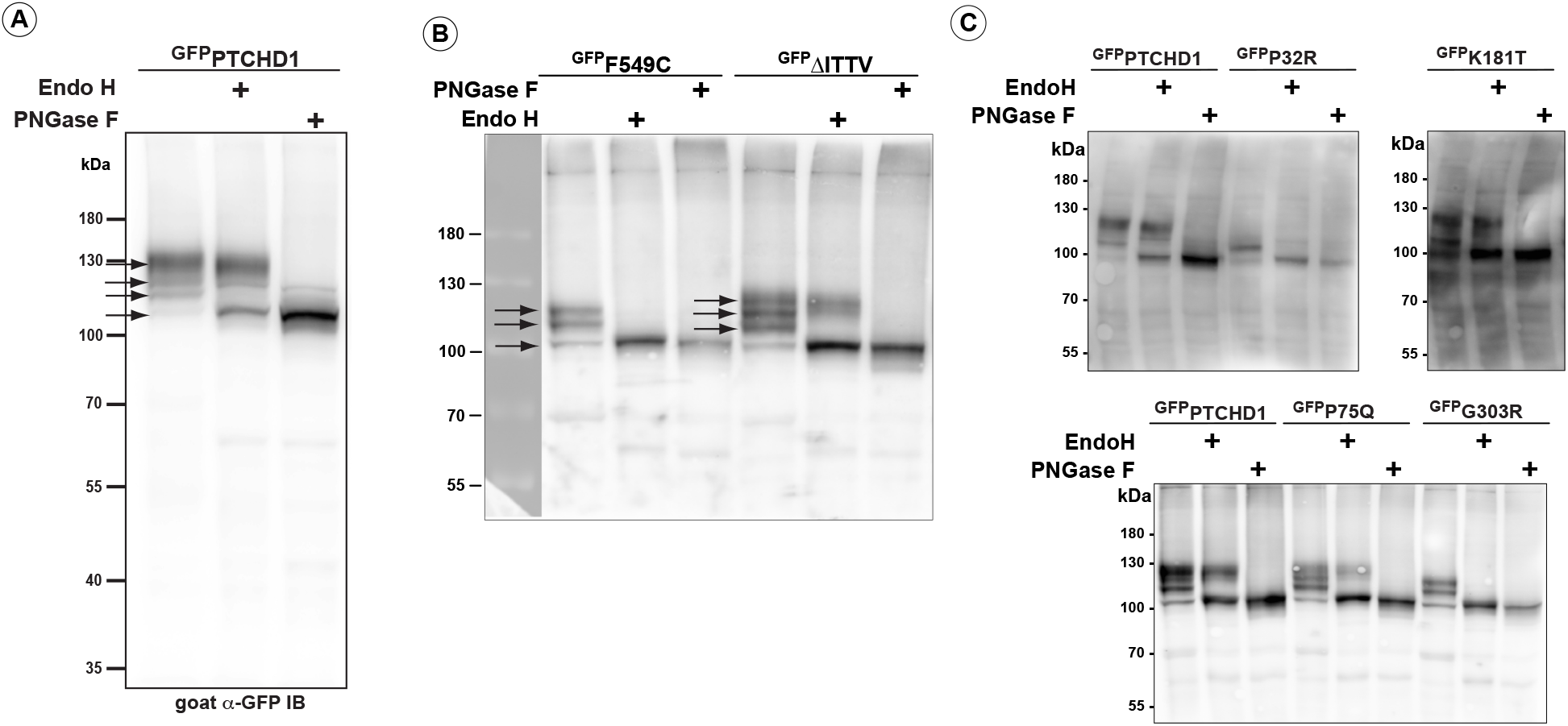
Altered post-translational processing of distinct PTCHD1 mutants. HEK293T cells were transfected with constructs expressing wild-type ^GFP^PTCHD1 or the point mutants, treated EndoH or PNGase and resolved by SDS_PAGE on western blots. **A)** Wild-type ^GFP^PTCHD1 alone. **B)** ^GFP^F549C (*left side*) and ^GFP^ΔITTV (*right side*) are differentially processed, only ^GFP^ΔITTV have the apparent normal processing. **C)** Analysis of processing for ^GFP^P32R, ^GFP^K181T, ^GFP^P75Q and ^GFP^G303R.

The fidelity of glycosylation plays a crucial role in stabilizing the protein expression in the cell. Given that specific point mutations modify *N*-linked glycosylation of ^GFP^PTCHD1, the stability of these mutant proteins were measured to determine the lack of maturation of glycosylated mutants correlated with altered protein half-life. As shown in **Figure 4A**, WT ^GFP^PTCHD1 protein exhibited a half-life beyond 12h, determined in HEK293 cells transiently expressing ^GFP^PTCHD1 and treated with cycloheximide (CHX). Similarly, the point mutant, P75Q exhibited similar stability relative to WT ^GFP^PTCHD1, consistent with its apparent normal processing. In contrast, the protein half-life of P32R, which exhibited immature processing, was decreased to 2.5h. **Figure 4B** shows that the concordance between processing and protein half-life is evident for the mutants K181T, F549C and the C-terminal truncation, ΔITTV. Curiously, the G303R mutant, which also failed to be processed to more mature forms, had a half-life similar to the wild-type protein.

**Figure 4.**
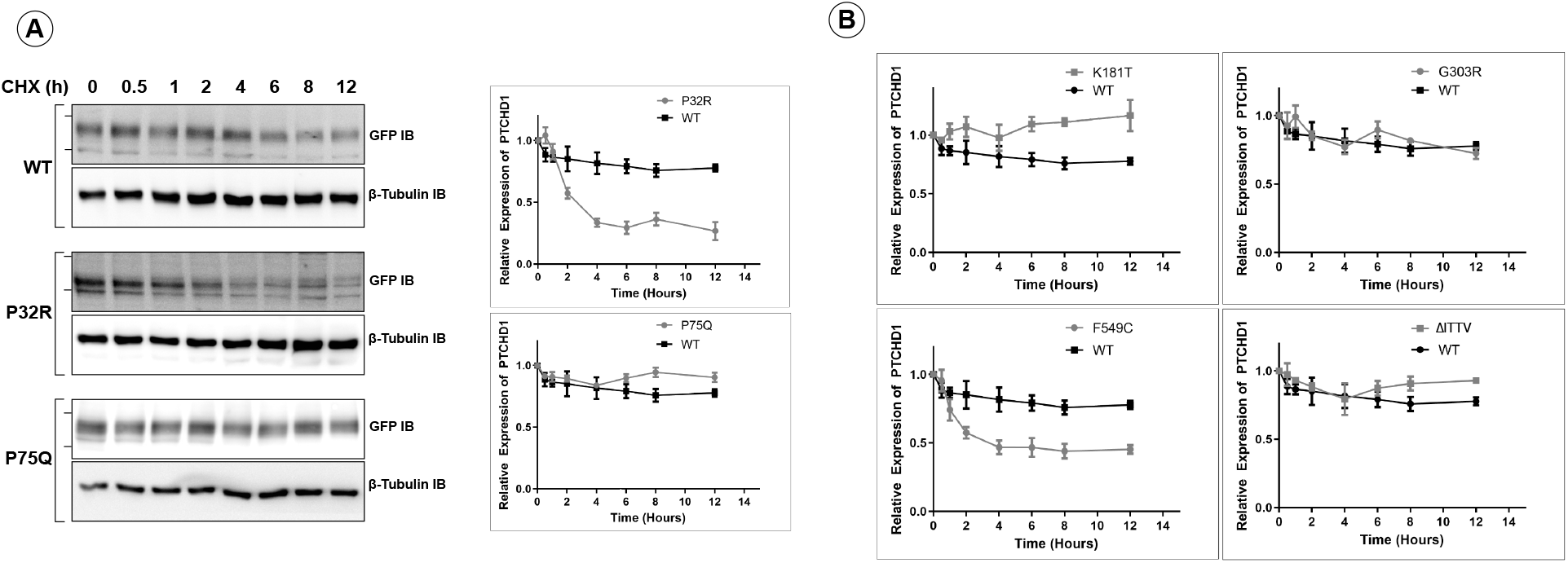
Point mutants alter the proteins stability of the P32R and F549C. HEK293T cells were transfected with one of the ^GFP^PTCHD1 point mutants and treated with cycloheximide at 50μg/mL for the time points indicated. **A)** *Left side* - Representative western blots for time course of CHX-treated HEK293 cells transiently expressing WT ^GFP^PTCHD1, ^GFP^P32R or ^GFP^P75Q mutants. *Right side* - Quantification of westerns blots. Relative levels of ^GFP^P32R or ^GFP^P75Q is compared to ^GFP^PTCHD1. **B)** Relative levels of ^FP^F549C, ^GFP^ΔITTV, ^GFP^K181T, and ^GFP^G303R. Error bars represent standard error of the mean value of at least 3 independent experiments.

Proper post-translational modifications play a fundamental role in subcellular targeting. Due to the defects of processing in ^GFP^PTCHD1 mutants harbouring specific point mutations, the subcellular localization may be affected. Specifically, proteins not susceptible to full EndoH cleavage may experience aggregation in the ER or Golgi, preventing their proper localization. We first tested the localization of the various mutants in transient assay in HEK293T cells. As **Figure 5** illustrates, no apparent alteration in the distribution of these mutants was evident. Here, signal for each of the GFP-tagged proteins was seen throughout the cytoplasm. There was no apparent defect in transport from the ER nor did any of the mutants exhibit localization to the Golgi, indicated by the lack of ^GFP^PTCHD1 signal that coincided with the Golgi marker, GM130.

**Figure 5.**
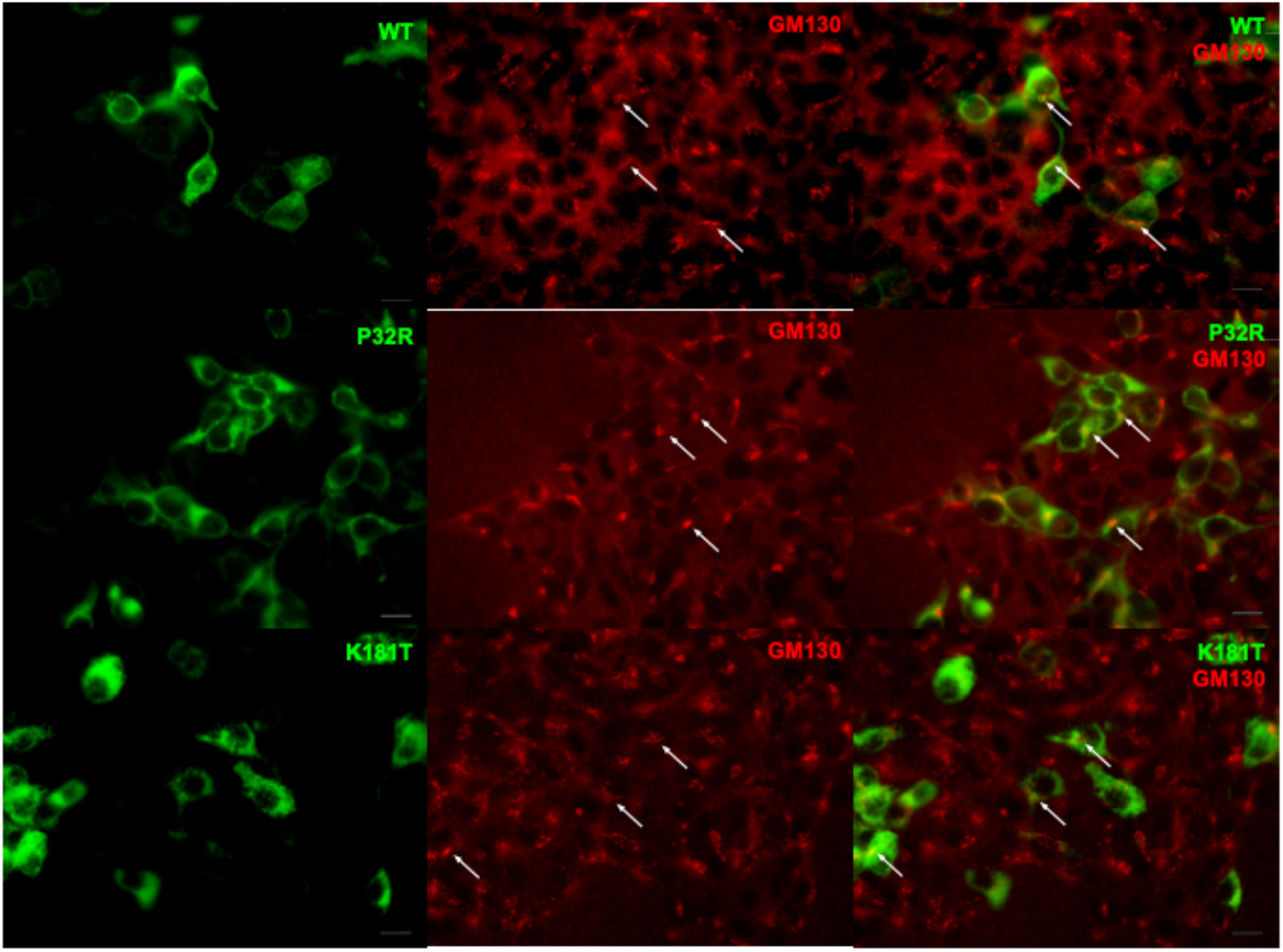
Transiently expressed PTCHD1 mutants in HEK293T cells show no apparent alteration in protein distribution. Immunofluorescence was performed using HEK293T cells transfected with ^GFP^PTCHD1, ^GFP^P32R or ^GFP^K181T (Green) and co-stained with an endogenous Golgi marker, GM130 (Red). All mutants localized similar to wild-type protein and showed no accumulation in the ER or golgi.

Previous studies showed ^GFP^PTCHD1 localizes with PSD95 in neuronal processes (18). As **Figure 6** shows for primary neurons, lentiviral-mediated expression of the P75Q, K181T, and P32R mutants all retained their ability to co-localize with PSD95 in neuronal processes. Thus, regardless of their processing and stability, these mutants do not exhibit altered intracellular localization.

**Figure 6.**
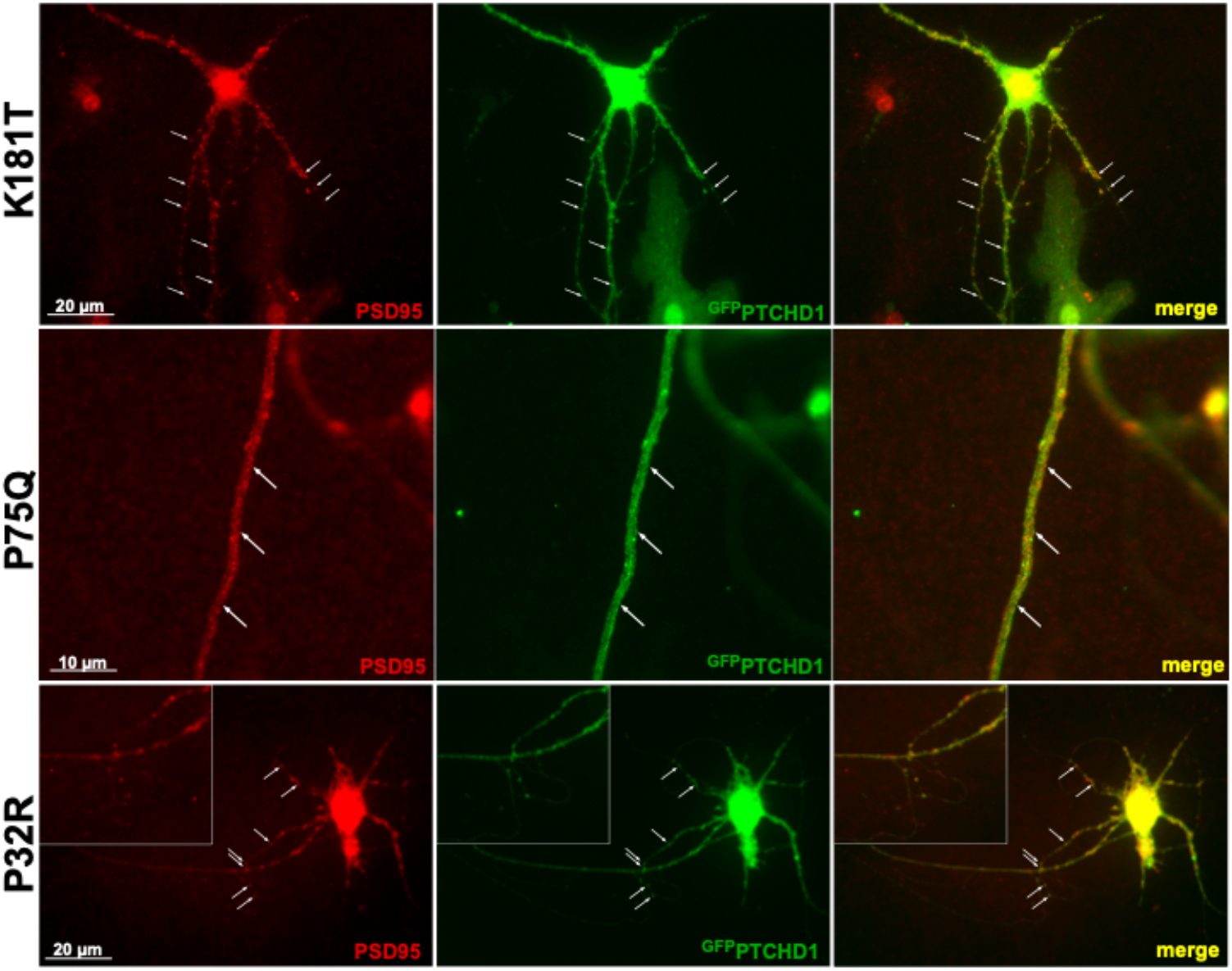
PTCHD1 constructs localize to endogenous PSD95 in primary neurons. Immunofluorescence was performed in primary neurons transiently-expressing ^GFP^PTCHD1 mutants, ^GFP^P32R, ^GFP^P75Q or ^GFP^K181T (green) co-localize with endogenous PSD95 (Red).

The isolated C-terminal domain of PTCHD1 was shown previously to bind to the PDZ-containing, postsynaptic scaffolding proteins, PSD95 and SAP102 (18). Using SAP102b^myc^, an isoform that only contains the third PDZ domain, as well as the SH3 and GK domains (34, 35), and the mouse ortholog of ^3xFlag^Ptchd1, **Figure 7** shows that all of the WT ^3xFlag^Ptchd1, as well as for the P75Q, G303R, F549C, and P32R co-immunoprecipitated SAP102b^myc^. That this interaction was mediated by the PDZ-binding motif at the very end of the C-terminus of Ptchd1 was confirmed using the deletion mutant, ΔITTV which harbours a deletion of the PDZ-binding motif.

**Figure 7.**
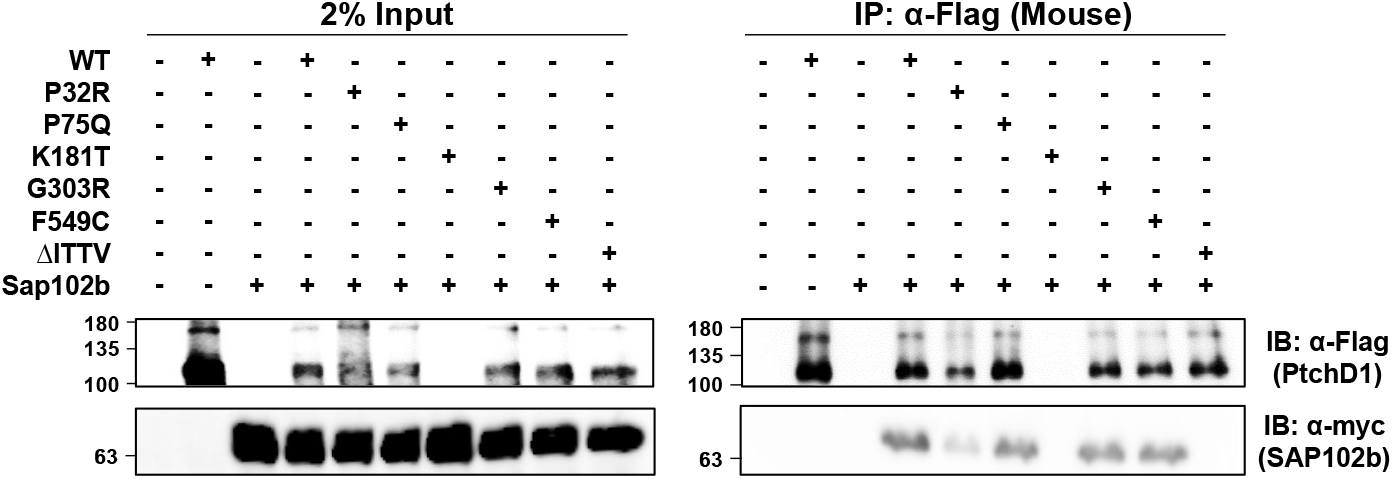
PTCHD1 interacts and co-localizes with SAP102b. *Left Side* - HEK293T cells were transiently transfected with either ^3xFlag^Ptchd1 or one of the 3xFlag-tagged point mutants (P32R, P75Q, G303R, F549C, or ΔITTV) and SAP102b^myc^, and resolved by SDS-PAGE on western blots. *Right Side* –Co-immunoprecipitation of SAP102b^myc^ with wildtype ^3xFlag^Ptchd1 or Ptchd1 point mutants. Each IP was divided in half, run on two gels and probed with either anti-flag (Ptchd1; upper panel) or anti-myc (SAP102b; lower panel)

Taken together our data suggest that the PTCHD1 missense mutants under investigation may exert an etiopathogenic effect through reduced lifespan of the protein and thus reduced bioavailability, rather than through disruption of the transport of PTCHD1 to its functional destination.

## Discussion

There exists a high density of synaptic scaffolding proteins at the PSD that organize neurotransmitter receptors, in part, by utilizing their PDZ domains to bind to cellular elements. Previously, interactions between the C-terminal PDZ binding motif in PTCHD1 with synaptic scaffolding proteins, PSD95 and SAP102 were described (17, 18). However, neither of these previous studies verified the interaction with the full-length PTCHD1 protein. We showed the interaction between full length PTCHD1 and SAP102. Further, using the shorter construct of SAP102b isoform, we demonstrated that the third PDZ domain of SAP102 was sufficient for PTCHD1 binding. SAP102 links NMDARs to excitatory type 1 synapses and mediates AMPAR-regulatory activities (34, 36, 37). This highly mobile member of the MAGUK family is not sequestered at the PSD as it has been shown to interact with NMDAR subunits in the secretory pathway, specifically the ER (35, 38). SAP102 also mediates NMDAR exocytosis, and activity distinct from those mediated by PSD95 (39). Truncating mutations of *DLG3*, the gene encoding SAP102, have been associated with X-linked retardation (40, 41) and XLID with substantial impairment in cognitive abilities and social and behavioural adaptive skills (42). Therefore, SAP102 has been proposed to be a plausible candidate gene for ASD (43). Its interaction with PTCHD1 may implicate that the mechanism employed for PSD-targeted trafficking and endocytosis of PTCHD1 is mediated by SAP102. In this vein, we have shown that all Ptchd1 point mutants studied here, except for ΔITTV which lacks the four-amino acid (ITTV) putative PDZ-binding domain, possess the ability to bind to SAP102b. These data suggest that the neurodevelopmental consequences of these specific *PTCHD1* mutations may arise independently of its interaction with SAP102 within dendritic spines.

Advances in genome sequencing and genetic analyses have identified an increasing number of genes responsible for ASD by detecting *de novo* mutations linked to ASD. These mutations can be CNVs or single-base-pair mutations; however, missense mutations are less informative because their impact on the protein is unknown. ASD-linked single missense mutations have been described in proteins involved in the dendritic spines such as *SHANK3, NLGN4*, and *ACTN4* (44–47). Single missense mutations have also been identified in *PTCHD1* from individuals with ASD or ASD-associated disorders. In this study we have analyzed a number of clinically-relevant PTCHD1 point mutations and found that P32R, G303R and F549C have the most adverse effects on the protein. These mutations affect *N*-linked glycosylation in post-translational modifications and with some resulting in protein destabilization. However, these mutants were not observed to cause aggregation in the ER.

The deleterious consequences on processing and stability that arise from the P32R and G303R mutations, respectively, are of particular interest. While the specific activities of PTCHD1 in dendritic spines remains unresolved, the primary and 3D structures of PTCHD1 suggest that it may harbor cholesterol transport activities similar to the related proteins, NPC1 (12, 13, 48–50) and Ptch1. Interestingly, both P32R and G303R in PTCHD1 are in similar positions to the residues in Ptch1 that were recently shown to be involved in an apparent complex with a cholesterol moiety observed in the cryo-EM of Ptch1 (22, 51). Located on the a-helices, TM1 and TM3, respectively, the sidechains of these residues are involved in coordinating the orientation of a likely cholesterol molecule in this region. Given the fundamental roles of cholesterol in the localization and activities of the synaptic transmitters (52–56), we suggest that these mutations in PTCHD1 may specifically alter synaptic signaling due to impaired localized transport of cholesterol in dendritic spines.

When observing the extent of maturation of *N*-linked glycosylation in PTCHD1 mutants, we hypothesized that the inherent stability of PTCHD1 could be altered due to an inability to be properly glycosylated. Proteins with post-translational modifications that are not properly matured do not typically proceed through the ER and Golgi and are subjected to early degradation. Our protein stability time course assay showed only two mutants, P32R and F549C, with a shorter half-life than WT. Both mutants exhibited some processing, these forms being susceptible to cleavage by Endo H indicating that they were not processed to the more mature Endo H-resistant forms. These results illustrate that single amino acid substitutions may result in decreased PTCHD1 protein stability as immature stages of *N*-linked glycosylation may not achieve proper folding of the protein and may lead to early degradation of the protein. The significance of this decreased stability requires further analysis.

In conventional protein processing pathways, alterations in glycosylation may result in aggregation in the ER, causing ER stress and preventing the trafficking to subcellular locations. Immunofluorescence staining of these PTCHD1 mutants showed none of the mutants were aggregated adjacent to the co-stain of a Golgi marker, GM130. Indeed, staining of PTCHD1 mutants was seen throughout the cell with no observable differences in localization between the WT PTCHD1 protein and the mutants. Likewise, regardless of whether mutants were fully processed, they were observed to co-localize with PSD95 in neuronal processes. We propose that these mutants may represent hypomorphic or null variants whose principal activities in the PSD are crippled or lost, respectively, despite their ability to localize to the PSD95-containing dendritic structures. While potentially mediating localized fluxes of cholesterol seems possible, analogous to the activities of the structurally-related proteins, NPC-1 and Ptch1, the uncharacterized activities of PTCHD1 remain speculative.

## Experimental Methods

### Cell Culture

Human embryonic kidney 293T (HEK-293T) cells (a kind gift from Prof. S. Girardin) were cultured in Dulbecco’s Modified Eagle Medium with 10% fetal bovine serum (FBS) (Wisent) and 1% penicillinstreptomycin (Wisent).

### Primary Neurons

Dissociated cortical neurons were prepared as described (57). In brief, cortical layer was dissected out of P0-P2 C57 and dissociated enzymatically (papain, 12 U/mL; Worthington) and mechanically (trituration through flame-polished Pasteur pipette). After dissociation, the cells were washed, centrifuged and plated on poly-d-lysine-coated glass coverslips at a density of 1.25-2.5E5 cells/mL. Growth media consisted of Neurobasal and B27 (50:1), supplemented with penicillin-streptomycin (50 U/mL; 50 U/mL) and 0.5 mM L-glutamax (Gibco). FBS (2%; Wisent) was added at the time of plating. After 5 d, half of the media was changed without serum and with cytosine arabinoside (5μM; Sigma-Aldrich) to limit proliferation of non-neuronal cells. Twice a week thereon, half of the growth medium was replaced with serum- and cytosine arabinoside-free medium.

### Construct Creation

The GFP-PTCHD1 in pDEST-53 vector construct was used to generate the panel of PTCHD1 point mutants. All single nucleotide substitution point mutations in the N-term GFP form were created with ligation-independent PCR cloning technique. Primers for the sense strand and anti-sense strand were designed for each independent mutation as seen in Appendix 2). The new construct was created from a PCR reaction that amplified the entire plasmid harbouring the substitution using the high-fidelity polymerase Q5 according to the manufacturer’s instructions (New England Biolabs; NEB). The unpurified PCR product was then treated with DpnI to digest the methylated template DNA which was subsequently transformed into DH5α. Final constructs were verified by Sanger sequencing (ACGT Corporation, Toronto). The SAP102 construct was kindly provided by Prof. Igor Stagljar (University of Toronto).

All PTCHD1 WT and point mutant constructs were inserted into the 3rd generation lentivirus vector, pUltra (Addgene #24129), which includes a puromycin resistance gene. Using PCR amplification, a unique NheI site was generated upstream of the start codon of the PTCHD1 constructs with a blunt end after the stop codon. The PCR product was ligated into the NheI and HincII sites in the pUltra vector. All clones were verified with restriction enzyme diagnostic digest.

The mouse ortholog of *Ptchd1*, which exhibits 98.1% sequence conservation (871 of 888 amino acids) with human *PTCHD1*, was amplified using high-fidelity Q5 polymerase from cDNA that was generated from RNA obtained from P19-induced neural cells. PCR amplicons were digested and ligated into pcDNA3.1 myc-His B expression vectors (Invitrogen), with a 3xFlag epitope tag also fused to the N-terminus of *Ptchd1*. Site-directed mutagenesis was subsequently used, as previously described, to generate the P32R, P75Q, K181T, G303R, F549C and ΔITTV point mutants of Ptchd1. Final constructs were verified by Sanger sequencing (The Centre for Applied Genomics, Toronto). Primer sequences for cloning and *Ptchd1* site-directed mutagenesis are provided in Appendix 2B.

### Lentivirus Production and Transduction

HEK293T cells were seeded in a 6cm plate. At ~80% confluency, transfection was performed with PEI (2 mg/mL) at a 2 μL:1 μg ratio of PEI to DNA in serum-free Dulbecco’s Modified Eagle Medium. Three plasmids were transfected: 1 μg of pLenti-PTCHD1, 0.75 μg psPAX2 packaging plasmid (Addgene #12260), and 0.25 pCMV-VSV-G envelope plasmid (Addgene #8454). After a 15 min incubation with PEI, the solution was added to the cells. Media was collected after 48 h and stored in 1 mL aliquots at −80°C.

Primary neurons were used for lentiviral transduction at D3. Transduction of cells were performed at 1:5 – 1:2 of viral media to total culture media and incubated for 48 h.

### Western Blotting and Co-immunoprecipitations

All western blots and co-immunoprecipitations were performed using lysates from transiently transfected HEK293T cells, unless stated otherwise. HEK293T cells were grown to 70-80% confluency in 100mm plates and transfected using 2mg/mL polyethylenimine (PEI, Sigma) at a 2μL:1μg ratio of PEI to DNA. Cell lysates were taken 48-72h after transfection. Cell lysates were prepared by washing cells twice with ice cold PBS, pH 7.4 (137mM NaCl, 2mL KCl, 10mM Na_2_HPO_4_, 2mM KH_2_PO_4_) then adding 1% NP-40 lysis buffer (50mM Tris pH 8.0, 120mM NaCl, 1% NP-40) containing protease and phosphatase inhibitors (10mM NaF, 1mM PMSF, 2μg/mL leupeptin, 2μg/mL aprotinin, 1mM sodium orthovanadate). For western analysis of protein expression, 50μg of lysate were used and samples in 4x-sample buffer were then incubated at 37°C for at least 20min prior to loading on the gel in order to avoid boiling-induced aggregation of SSD-containing transmembrane proteins (14, 58). Blots were developed using Western Lightning PLUS ECL (PerkinElmer) and imaged on a MicroChemi 2.0 Imager (FroggaBio). Antibodies and concentrations used for western blots can be seen in Appendix 3.

Immunoprecipitation experiments were performed using 250μg of total protein, made up to 500μL total volume with 1% NP-40 buffer. Samples were incubated overnight at 4°C with primary antibody. Immuncomplexes were bound to Protein G-Agarose (Invitrogen) or Protein A-Agarose (Pierce) beads and washed 5x 1% NP-40 buffer and blots prepared as described above.

For co-immunoprecipitation experiments involving ^3xFlag^Ptchd1 and SAP102b^myc^, 250 μg of total protein samples were made up to 500 μl total volume with 1% NP-40 buffer and incubated overnight at 4°C with 1 μl of mouse anti-Flag primary antibody (#F1804; Sigma). Subsequently, samples were incubated with 50 μl of Protein G-conjugated Dynabeads (#10003D; Invitrogen) for two hours at room temperature, and immune complexes were eluted (75 mM Glycine-HCl, pH 2.7) for five minutes at room temperature. For western blot detection of ^3xFlag^Ptchd1 and SAP102b^myc^, rabbit anti-DYKDDDDK (#D6W5B; NEB) and rabbit anti-myc (#71D10; NEB) primary antibodies were used, respectively, followed by the anti-rabbit HRP-conjugated secondary antibody (#W4011; Promega).

### Glycosylation Assay

N-linked glycosylation processing of PTCHD1 was determined by Endo-β-N-acetylglucosaminidase H (Endo H) and Peptide-N^4^-(N-acetyl-β-glucosaminyl)-asparagine amidase F (PNGase F) as we previously described (14). Briefly, PTCHD1 mutants were transiently expressed in HEK293T cells. Cells lysed in 1% NP40 buffer and 50μg of protein lysate was made up to a total volume of 20μL containing ddH_2_O and the appropriate NEB enzyme buffers. The samples were treated with either **i)** no enzyme, **ii)** 500U Endo H (NEB), or **iii)** 500U PNGase F (NEB) for 1h at 37°C. Proteins were resolved using SDS-PAGE and migration of the samples determined by western blot analysis.

### Protein Stability Assay

To determine the stability of PTCHD1 mutant proteins, HEK293T cells transiently expressing these proteins were treated with the 50μg/ml of the protein synthesis inhibitor, cycloheximide (CHX) for the described lengths of time. Cell lysates were prepared as described previously and the expression levels of PTCHD1 was determined by western blot analysis. Relative expression of PTCHD1 protein was quantified by densitometry using ImageJ software. PTCHD1 signal was normalized to β-tubulin. Independent experiments were performed at least 3 times per construct. Statistical analysis was performed on GraphPad Prism.

## Acknowledgements

This work was supported by grants from the Canadian Institutes of Health Research to J.B.V. (MOP-114952 and #PJT-156367)

## Abbreviations

ASD: Autism spectrum disorder
CHX: cycloheximide
PEI: polyethylenimine
RA: retinoic acid
RND: resistance nodulation division
SSD: sterol-sensing domain
TM: transmembrane
XLID: X-linked intellectual disability.

## Appendix 1: selected PtchD1 variants for functional analysis

Appendix 1: selected PtchD1 variants for functional analysis.
| **Mutation** | Genomic (hg19);<br>cDNA (NM_173495.2)<br>coordinates | Source,<br>subject ID | Inheritance | Minor allele<br>frequency<br>(gnomAD)/#<br>hemizygotes | Prediction: Condel; CADD<br>PHRED-like scaled C-score | Reported phenotype |
| --- | --- | --- | --- | --- | --- | --- |
| P32R | ChrX:23353087;<br>c.95C>G | DECIPHER:<br>284363 | Mat | 0/0 | Deleterious; 26.8 | Autistic behavior; cognitive<br>impairment |
| P75Q | ChrX:23353216;<br>c.224C>A | MSSNG:<br>AU3794302 | Mat | 0/0 | Deleterious; 27.6 | Autism spectrum disorder |
| K181T | ChrX:23397898;<br>c.542A>C | Clinvar | nr | 0/0 | Deleterious; 21.9 | Abnormality of brain<br>morphology |
| G303R | ChrX:23398263;<br>c.907G>A | ClinVar ID<br>417957 | nr | 0/0 | Deleterious; 27.7 | Autism |
| F549C | ChrX:23411281;<br>c.1646T>G | Ptchd1-base.com | nr | 0/0 | Deleterious; 28 | Autism spectrum disorder |
| ΔITTV | ChrX:23412285;<br>c.2650C>T | No subject | na | 0/0 | 38 | na |

nr=not reported
na=not applicable

## Appendix 2. Primers used to generate PTCHD1 point mutations

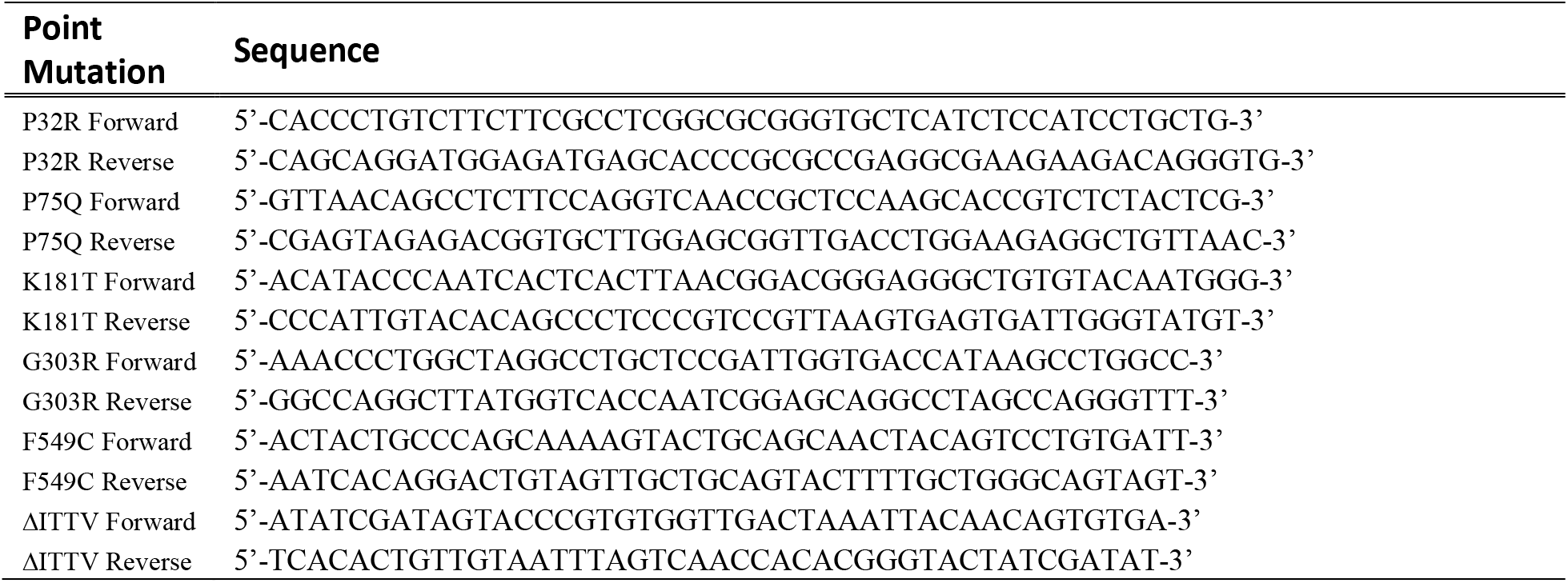

## Appendix 2B. Primers used to for Mouse *Ptchd1* cloning and site-directed mutagenesis

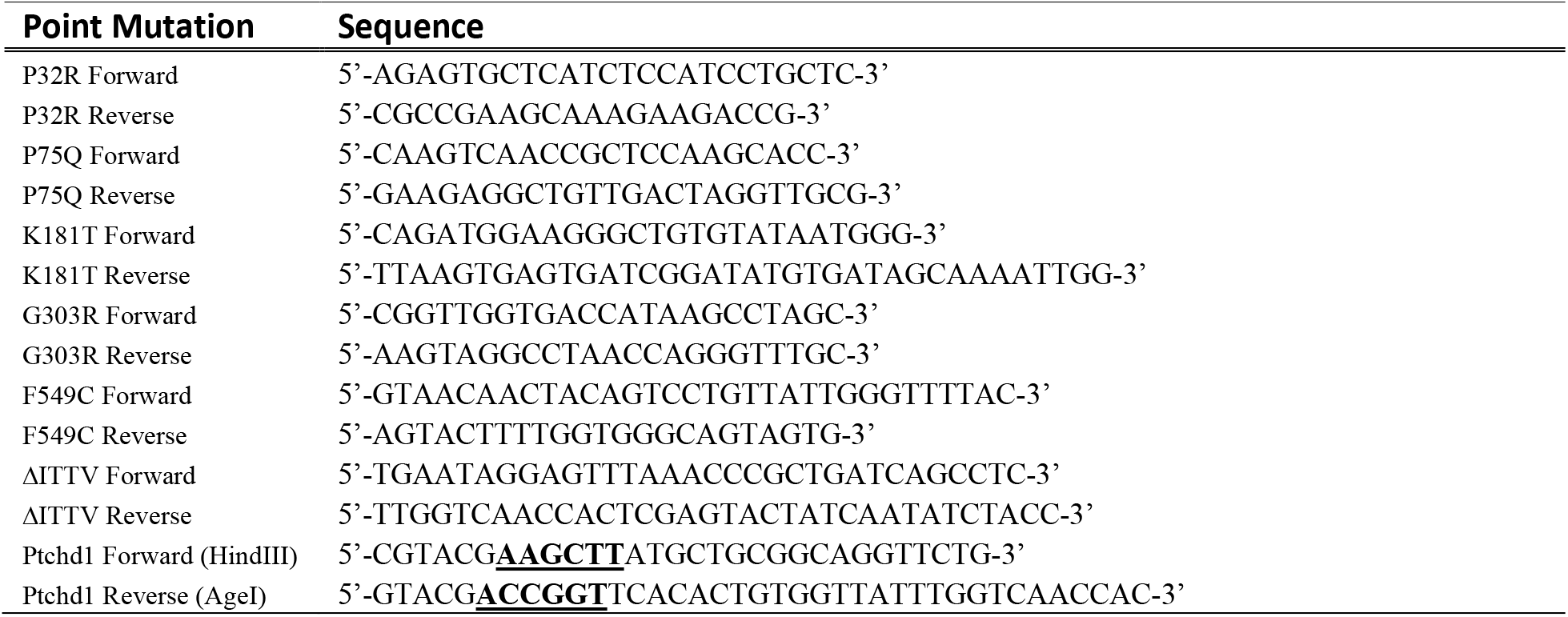

## Appendix 3. Antibodies used for western blots and immunofluorescence staining

Appendix 3. Antibodies used for western blots and immunofluorescence staining.
|  | Antibody | Source | Dilution |
| --- | --- | --- | --- |
| 1° Antibody | Mouse α-Flag-Tag | Applied Biological Materials Inc. Cat # G191 | 1:1000 WB IB |
| | Mouse α-Myc-Tag (9B11) | Cell Signaling Technologies #2276S | 1:1000 WB IB;<br>1:400 IF |
| | Rabbit α-GFP-Tag | Invitrogen Cat #A11122 | 1:100 WB IB;<br>1:400 IF |
| | Goat α-GFP-Tag | Applied Biological Materials Inc. Cat # G096 | 1:1000 WB IB;<br>1:400 IF |
| | Goat α-Biotin | Cedarlane, CLAS10-666) | 1:1000 WB IB |
| | Rabbit α-PSD95 | Abcam ab18258 | 1:400 IF |
| | Mouse α-β-Tubulin | Applied Biological Materials Inc. Cat # G098 | 1:5000 WB IB |
| | Mouse α-GM130 | BD Transduction Laboratories Cat # 51-9001978 | 1:400 IF |
| HRP-Linked 2° Antibody | Donkey α-goat IgG | Santa Cruz Biotechnologies Inc. sc-2020 | 1:5000 |
| | Horse α-mouse IgG | Cell Signaling Technologies #7076 | 1:5000 |
| | Goat α-rabbit IgG | Cell Signaling Technologies #7074 | 1:5000 |
| Immunofluorescence<br>2° Antibody | Donkey α-rabbit IgG Alexa Fluor 488 | Invitrogen A21206 | 1:400 IF |
| | Donkey α-mouse IgG Alexa Fluor 568 | Invitrogen A10037 | 1:400 IF |
| | Donkey α-rabbit IgG Alexa Fluor 568 | Invitrogen A10042 | 1:400 |
| | Donkey α-goat FITC | Jackson ImmunoResearch 705-095-003 | 1:400 |
|  | Phalloidin Alexa Fluor 350 | Invitrogen A22281 | 1:20 |

## Appendix 4. Single-cell RNAseq co-expression data extracted from adolescent mouse study (mousebrain.org; Zeisel et al, 2018), showing cell types with highest expression of *Ptchd1* (top 20, highest to lowest), with expression values (log2(x+1) transformed average molecule counts) for putative interactors, and for housekeeping genes

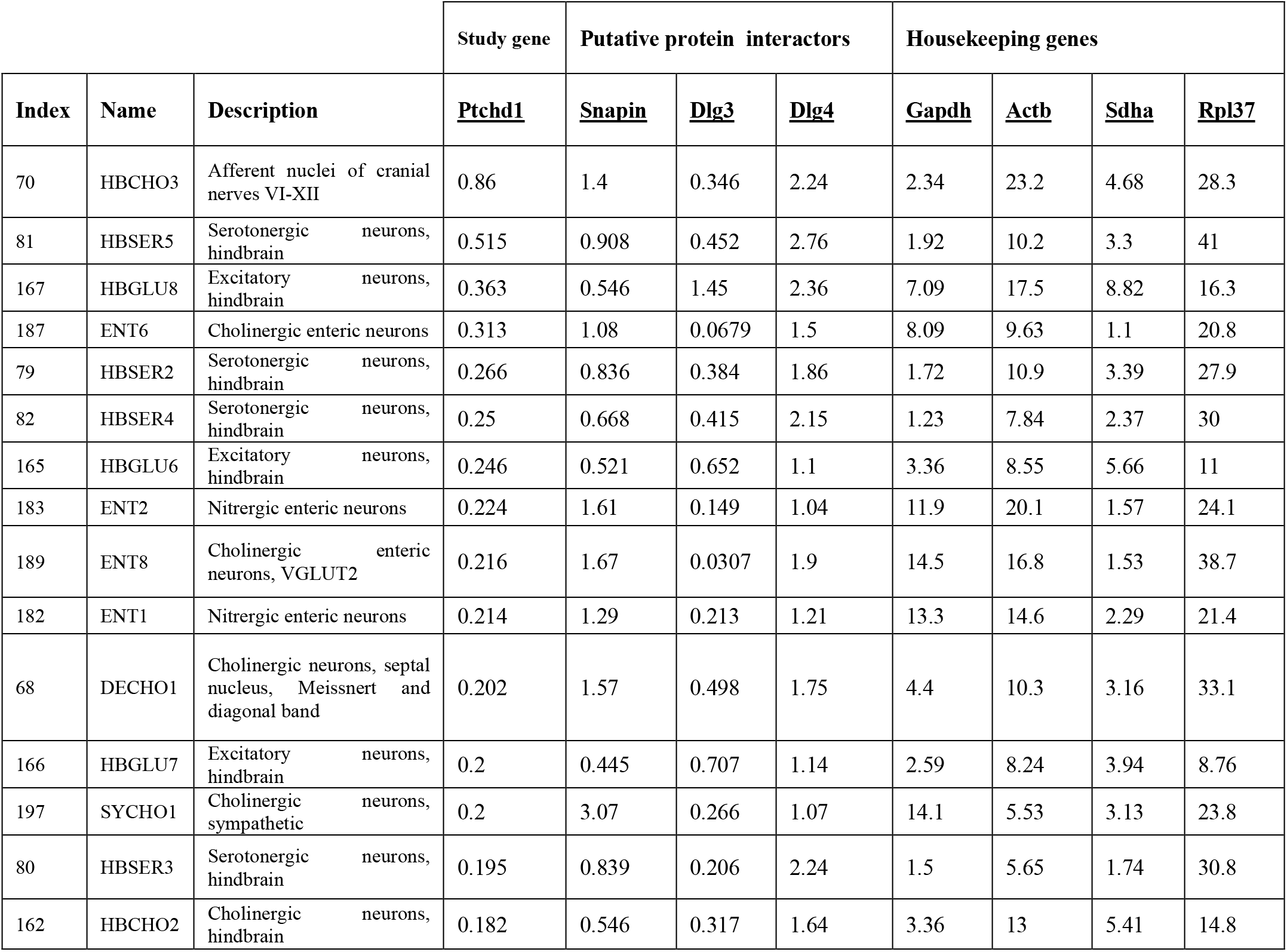
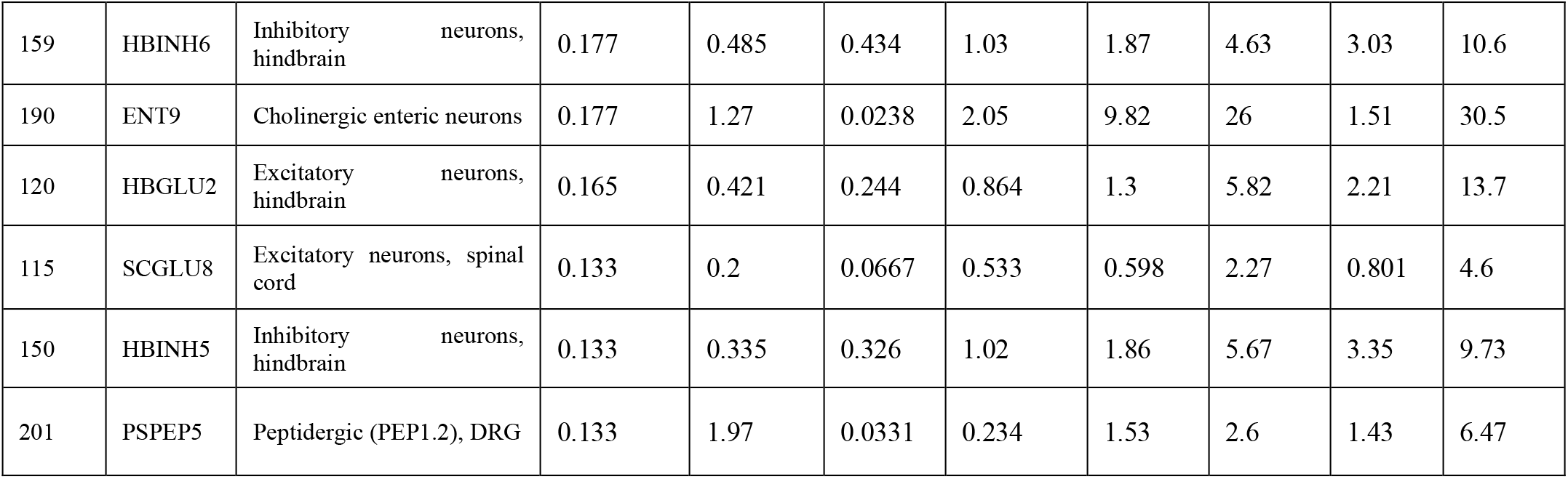

